# OpgH is an essential regulator of *Caulobacter* morphology

**DOI:** 10.1101/2023.08.28.555136

**Authors:** Allison K. Daitch, Erin D. Goley

## Abstract

Bacterial growth and division rely on intricate regulation of morphogenetic complexes to remodel the cell envelope without compromising envelope integrity. Significant progress has been made in recent years towards understanding the regulation of cell wall metabolic enzymes. However, other cell envelope components play a role in morphogenesis as well. Components required to maintain osmotic homeostasis are among these understudied envelope-associated enzymes that may contribute to cell morphology. A primary factor required to protect envelope integrity in low osmolarity environments is OpgH, the synthase of osmoregulated periplasmic glucans (OPGs). Here, we demonstrate that OpgH is essential in the ⍺-proteobacterium *Caulobacter crescentus*. Unexpectedly, depletion of OpgH results in striking asymmetric bulging and cell lysis, accompanied by misregulation of cell wall insertion and mislocalization of morphogenetic complexes. The enzymatic activity of OpgH is required for normal cell morphology as production of an OpgH mutant that disrupts a conserved glycosyltransferase motif phenocopies the depletion. Our data establish a surprising function for an OpgH homolog in morphogenesis and reveal an essential role of OpgH in maintaining proper cell morphology during normal growth and division in *Caulobacter*.

**Significance:** Bacteria must synthesize and fortify the cell envelope in a tightly regulated manner to orchestrate growth and adaptation. Osmoregulated periplasmic glucans (OPGs) are important, but poorly understood, constituents of Gram-negative cell envelopes that contribute to envelope integrity and protect against osmotic stress. Here, we determined that the OPG synthase OpgH plays a surprising, essential role in morphogenesis in *Caulobacter crescentus*. Loss of OpgH causes asymmetric cell bulging and lysis via misregulation of the localization and activity of morphogenetic complexes. Because cell envelope integrity is critical for bacterial survival, understanding how OpgH activity contributes to morphogenesis could aid in the development of antibiotic therapies.

## Introduction

Bacteria inhabit an impressive range of environments and can adapt to sudden changes in environmental conditions. One parameter that can vary significantly in different niches is osmolarity, ranging from dilute freshwater habitats to highly concentrated soil. The α-proteobacterium, *Caulobacter crescentus* (hereafter *Caulobacter*), is well-established as a model for physiological adaptation in the face of changing environments. Originally classified as an aquatic oligotroph, there is now evidence that *Caulobacter* also inhabits soil environments (1). Within these diverse habitats, *Caulobacter* is capable of exquisitely tuning its physiology as needed to propagate through growth and division.

The surrounding environment and available nutrients dictate *Caulobacter* cell cycle progression (2). *Caulobacter* undergoes distinct morphological changes as it proceeds through its cell cycle. A newborn, flagellated swarmer cell has the ability to move and search for nutrients before differentiating into a sessile stalked cell (2). This transition involves a stereotyped set of physiological changes including shedding its flagellum and growing a thin extension of the cell envelope called the stalk (2). The stalked cell can then undergo a cycle of cell division, as characterized by replication and segregation of the chromosome, elongation of the cell body, growth of a flagellum opposite the stalked pole, and cytokinesis (2). This asymmetric life cycle enables swarmer cells to find a new environment while requiring stalked cells to adapt within a given environment and any changes they experience there.

*Caulobacter* has distinct cellular structures and processes that allow for its survival in changing environments. Notably, the bacterial cell envelope serves as the physical barrier between the cell and its environment (3). The Gram-negative cell envelope comprises the inner and outer cell membranes and the periplasm between them. Within the periplasm is the peptidoglycan (PG) cell wall which provides structure and shape to the cell and protects the cell from lysis due to turgor pressure (4). PG biosynthesis is coordinated largely by the localization and activity of two conserved morphogenetic complexes: the elongasome/Rod complex and the divisome, responsible for elongation and division, respectively (5). Without the cell wall, significant reduction in osmolarity would quickly alter cell shape and result in lysis.

In addition to the cell wall, some bacteria also produce glucose polymers in the periplasm, called osmoregulated periplasmic glucans (OPGs, also referred to as membrane-derived oligosaccharides). In *Escherichia coli*, OPGs increase in abundance in low osmolarity environments and are thought to function as osmoprotectants (6). Theoretically, OPGs modulate the osmolarity of the periplasm to protect the cytoplasmic osmolarity from environmental changes. OPGs are produced across proteobacteria, though their structures can vary significantly, ranging from 5 to 24 glucose units in linear or cyclic configurations, and sometimes bearing modifications from phospholipids or intermediary metabolism (6). We recently identified the first OPG in *Caulobacter:* a cyclic hexamer of glucose (7).

The synthase of OPGs has been characterized in other organisms as the inner membrane protein, OpgH. In *E. coli*, OpgH is necessary for OPG production (8). Some organisms, including *E. coli*, encode additional *opg* genes, such as *opgG and opgD*, which are postulated to modify OPGs (6). Surprisingly, however, the only *opg* homolog encoded in *Caulobacter* is *opgH* (*CCNA_02097)*. Also notably, *Caulobacter opgH* is annotated as essential (9), unlike characterized *opgH* homologs in other organisms. OpgH and other OPG synthetic enzymes have been functionally implicated in osmoprotection, antibiotic resistance, motility, virulence, and symbiosis, but have not been reported to be essential for viability in any organism studied thus far (10).

In this work, we explore the role of OpgH in *Caulobacter* growth and morphogenesis. We demonstrate the essentiality of *opgH* in *Caulobacter* and discover striking morphological defects associated with OpgH depletion or loss of OpgH enzymatic activity. Our data reveal a novel role for an OpgH homolog, and OPGs, in maintaining cell morphology under unstressed growth conditions.

## Results

### OpgH is essential in *Caulobacter*

This study was initiated through our interest in identifying essential components of the cell envelope that contribute to morphogenesis. Transposon sequencing indicated that *opgH* (*CCNA_02097)*, encoding a putative inner membrane-associated glucan glycosyltransferase, was essential in *Caulobacter* (9). This was surprising because in *E. coli*, *opgH* is non-essential and deletion of *opgH* yields minimal defects in unstressed conditions (11). We therefore sought to validate the predicted essentiality of *opgH* in *Caulobacter*. Indeed, we were unable to generate a deletion of *opgH*. We were, however, able to make an OpgH depletion strain. This strain has a deletion of *opgH* at the native locus with a vanillate-inducible copy of *opgH* at the *vanA* locus (EG3421).

When grown with vanillate to induce *opgH* expression (+OpgH), cells looked morphologically wild type (WT) in both the complex media peptone yeast extract (PYE) and in defined minimal media (M2G) (Figure 1A t=0). With OpgH, cells grew comparably to WT *Caulobacter* in both liquid (measured by optical density) (Figure 1B) and solid media (measured by spot dilution) (Figure 1C). When OpgH was depleted for three hours in PYE without vanillate (-OpgH), cells exhibited a slight elongation and widening of the cell body. This phenotype was exacerbated during extended depletion, with cells showing prominent morphological defects at five and seven hours of depletion, including asymmetric bulging and cell lysis (Figure 1A). The bulging and lysis phenotype was also present for cells grown in M2G, but began later in the course of depletion, likely owing to the longer doubling time of *Caulobacter* in M2G (Figure 1A). In addition to morphological defects, prolonged depletion of OpgH quickly became lethal in PYE or M2G, as seen by growth curve (Figure 1B) and spot dilution (Figure 1C) assays. Slight overexpression of *opgH* from the vanillate-inducible promoter in the presence of native *opgH* did not impact cell growth (Figure 1B,C *vanA::P_van_-opgH*).

**Figure 1.**
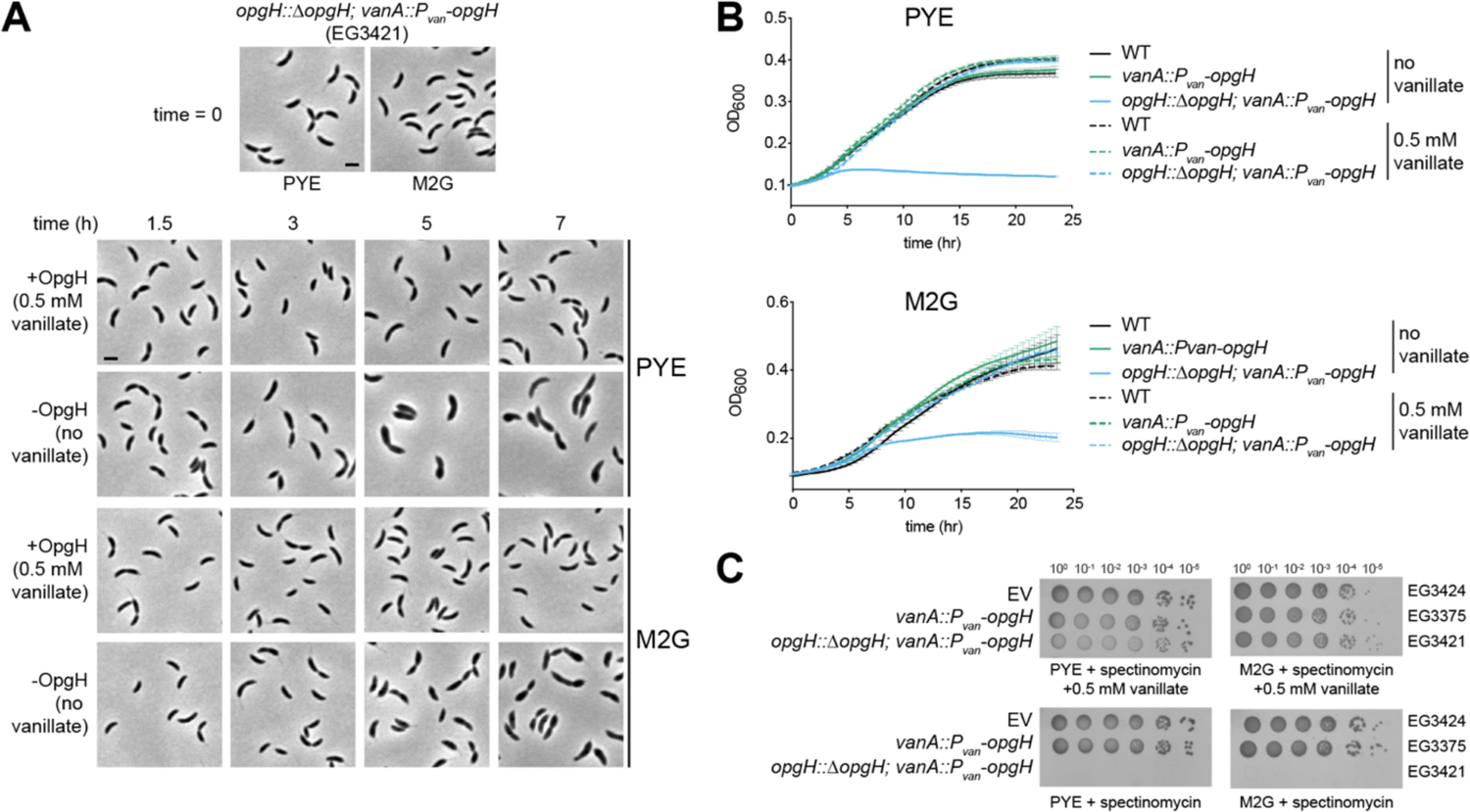
OpgH is essential for viability. **A.** Phase contrast images of OpgH depletion strain (EG3421) in the presence and absence of 0.5 mM vanillate over the course of seven hours. Cells were either grown in complex media (PYE) or minimal media (M2G) as indicated. Scale bar, 2 µm. **B.** Growth curve of WT (EG865), OpgH over-producing (EG3375), and OpgH depletion (EG3421) strains in PYE or M2G. **C.** Spot dilutions of empty vector (EV, EG3424), OpgH over-producing (EG3375), and OpgH depletion (EG3421) strains on PYE or M2G plates with indicated antibiotics and inducer.

We were surprised by the speed at which we began to see morphological defects from depleting OpgH. Notably, we saw morphological changes beginning around three hours of depletion, which is approximately two cell cycles in PYE. Typically, depletion of a protein requires growth to reduce the available pool of protein, and morphological changes require enzymatic remodeling of PG during growth and division. Since the morphological defects we observed occurred fairly quickly, we wondered whether the morphology change resulting from OpgH depletion required active cell growth. To test this, we depleted OpgH in the presence of sub-lethal concentrations of chloramphenicol to inhibit cell growth. Depleting OpgH in the absence of chloramphenicol yielded the expected asymmetric bulging and lysis, however, these phenotypes were suppressed in the presence of chloramphenicol (Supplemental Figure 1). This suggested that active growth is needed to deplete OpgH and/or enable morphological phenotypes and ultimate lethality.

### OpgH depletion causes morphological defects including asymmetric bulging

The unique bulging phenotype of the depletion observed by eye motivated us to quantify the shape changes resulting from loss of OpgH. Using CellTool (12) to perform principal component analysis of cell morphology, we analyzed the OpgH depletion strain after five hours in the presence or absence of vanillate in PYE. We saw a statistically significant difference between cells with OpgH present (+OpgH) compared to cells without OpgH (-OpgH) in four shape modes (Figure 2A). These shape modes approximately reflected the following features: shape mode 1, length; shape mode 2, curvature; shape mode 3, width; and shape mode 4, asymmetric bulging. Cells depleted of OpgH for five hours were typically longer, less curved, and wider than cells with OpgH (Figure 2A). For shape mode 4, which reflects asymmetric bulging, we have reported the absolute values, since the bulging can appear on either side of a cell outline. Our analysis indicates that cells producing OpgH rarely, if ever, exhibit asymmetric bulging while this is frequently observed in the OpgH-depleted condition (Figure 2A, B). We also measured the shape variance of the four shape modes when OpgH was depleted in M2G. The OpgH-depleted cells in M2G (M2G - OpgH) have statistically significant differences in shape modes 1 through 4 compared to cells with OpgH present (M2G + OpgH) (Supplemental Figure 2).

**Figure 2.**
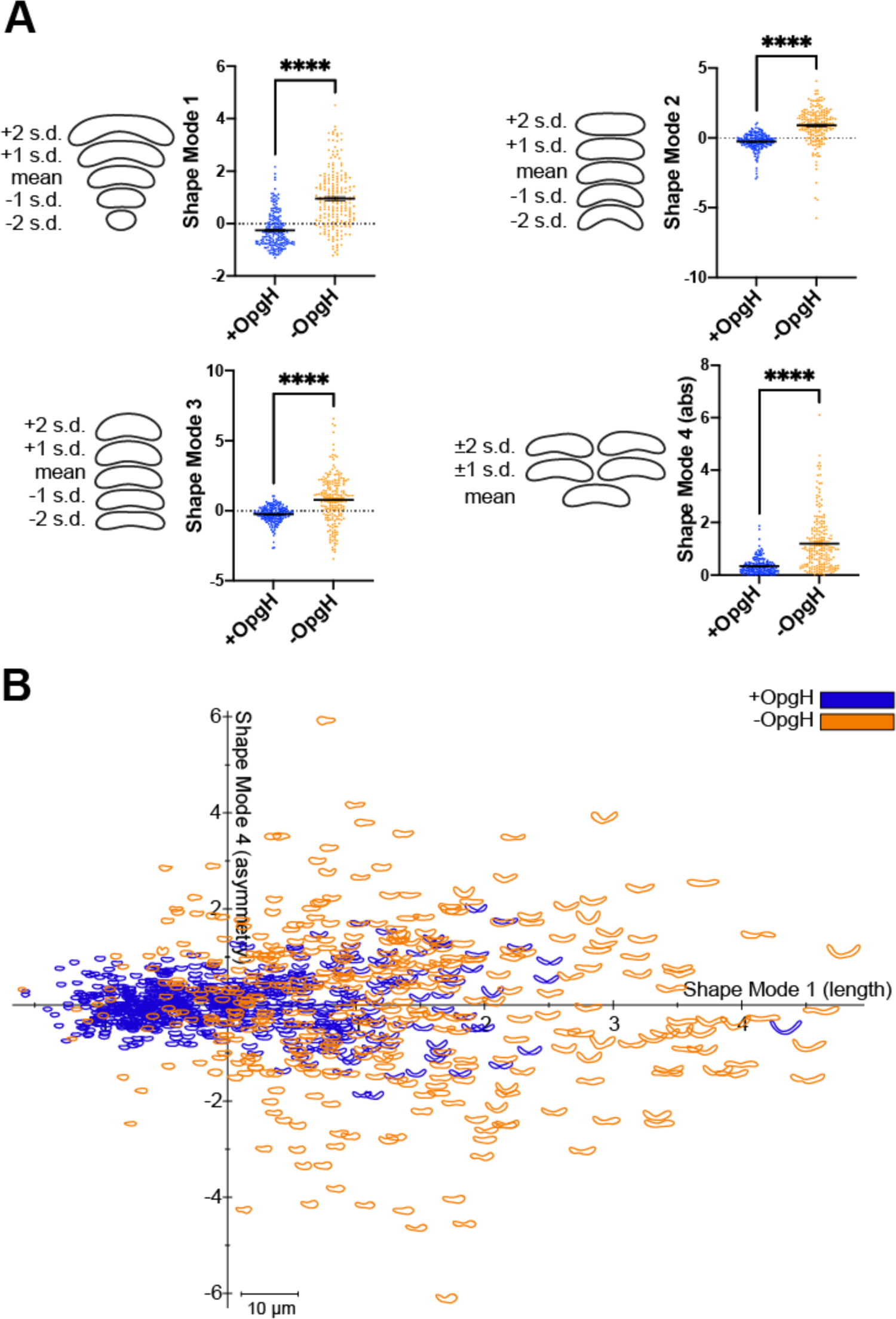
Cells lacking OpgH have morphological defects. **A.** Principal component analysis (PCA) of the OpgH depletion strain (EG3421) after 5 hours with 0.5 mM vanillate (blue, +OpgH) or without vanillate (orange, -OpgH). Scatter plots of 200 cells are shown for shape modes 1, 2, 3, and 4 which correspond to length, curvature, width, and asymmetric bulging. Contours are shown on the left to indicate the mean shape and 1 or 2 standard deviations from the mean for each shape mode. The absolute values are shown for shape mode 4. Statistical analysis uses a Mann-Whitney unpaired t-test. **** = P < 0.0001. **B.** Plot of shape mode 4 (asymmetric bulging) versus shape mode 1 (length) values for the OpgH depletion strain with vanillate (blue) or without vanillate for 5 hours (orange).

We were especially interested in the unique asymmetric bulging phenotype of OpgH-depleted cells. To investigate this asymmetry, we leveraged the inherent cell polarity of *Caulobacter* to determine if the bulging consistently occurred at the same pole or was random. Every *Caulobacter* cell has a defined orientation, with a stalked pole and a swarmer pole, that can be visually tracked over the course of a few cell cycles. We performed timelapse microscopy of the OpgH-depleted cells in the absence of vanillate to observe depletion over the course of five hours. The most notable phenotypes are summarized in Figure 3, including asymmetric bulging, elongation, multiple constriction events, and lysis. From our timelapse analysis, we were able to follow single cells through complete cell cycles, which allowed us to orient the cells and identify the stalked (old) or swarmer (new) pole of a given cell. As illustrated in the top three examples in Figure 3 (asymmetric bulges), we determined that bulging exclusively occurs in the stalked half of the cell (nearest to the old pole, indicated by black arrows). We localized an mNeonGreen fusion to OpgH and found that it was diffuse along the entire body of the cell in both PYE and M2G media (Supplemental Figure 3). These observations suggest that bulging is not related to OpgH localization, but to differential sensitivity of one or more factors unique to the stalked half of the cell during OpgH depletion.

**Figure 3.**
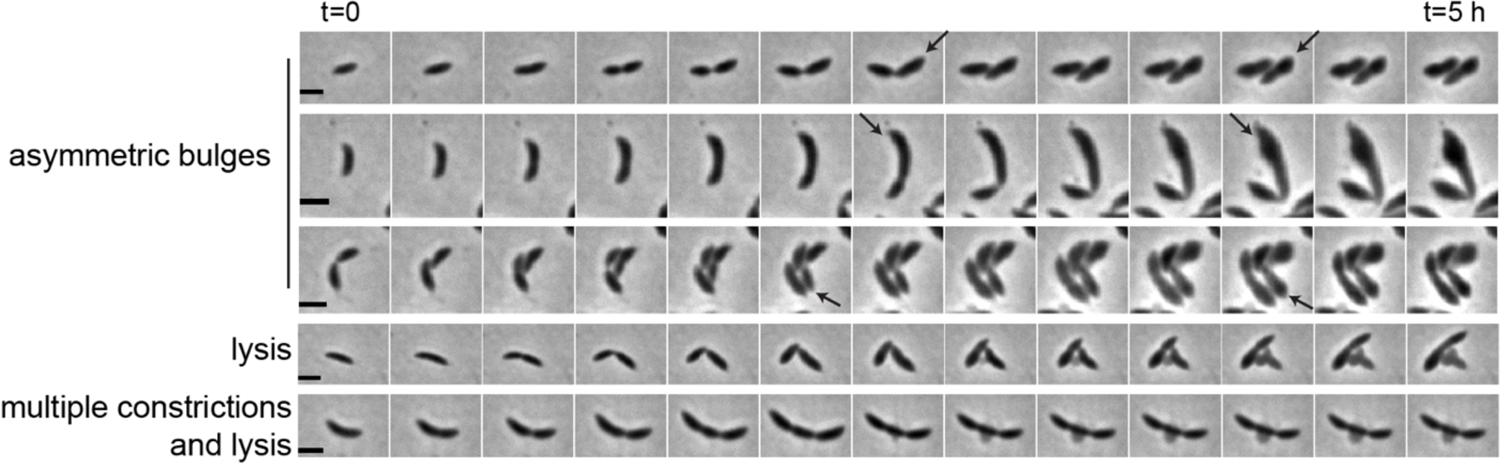
Depletion of OpgH causes bulging in the stalk-proximal region of the cell. Time-lapse micrographs of prominent phenotypes of OpgH depletion over the course of 5 hours (EG3421). Black arrows indicate old pole. Scale bar, 2 µm.

### Cell wall synthesis is disrupted during OpgH depletion

The primary determinant of cell shape is the PG cell wall, so we hypothesized that bulging is a result of misregulated PG synthesis. We therefore sought to visualize active PG synthesis in OpgH-depleted cells. Using the fluorescent D-amino acid, HADA (13), we captured a snapshot of active PG synthesis over the course of OpgH depletion. The HADA patterning of the OpgH depletion strain grown with vanillate (+OpgH) corresponded with the expected localization of PG synthesis in WT *Caulobacter*. HADA incorporated at the cell pole and/or broadly along the cell body in swarmer cells and then localized to mid-cell in stalked and pre-divisional cells (Figure 4A). This was consistent over the course of seven hours of growth with OpgH present. When vanillate was removed and OpgH was depleted (-OpgH), we began to observe atypical HADA incorporation starting at five hours (Figure 4B). At five hours of depletion, HADA incorporation was more diffuse, but still yielded a thick band near mid-cell that was typically adjacent to the asymmetric bulges and closer to the new cell pole than the old pole (Figure 4B). By seven hours of depletion, HADA incorporation was almost entirely diffuse and, in many cases, we observed minimal incorporation (Figure 4B). From these data we conclude that PG synthesis is spatially misregulated in the absence of OpgH.

**Figure 4.**
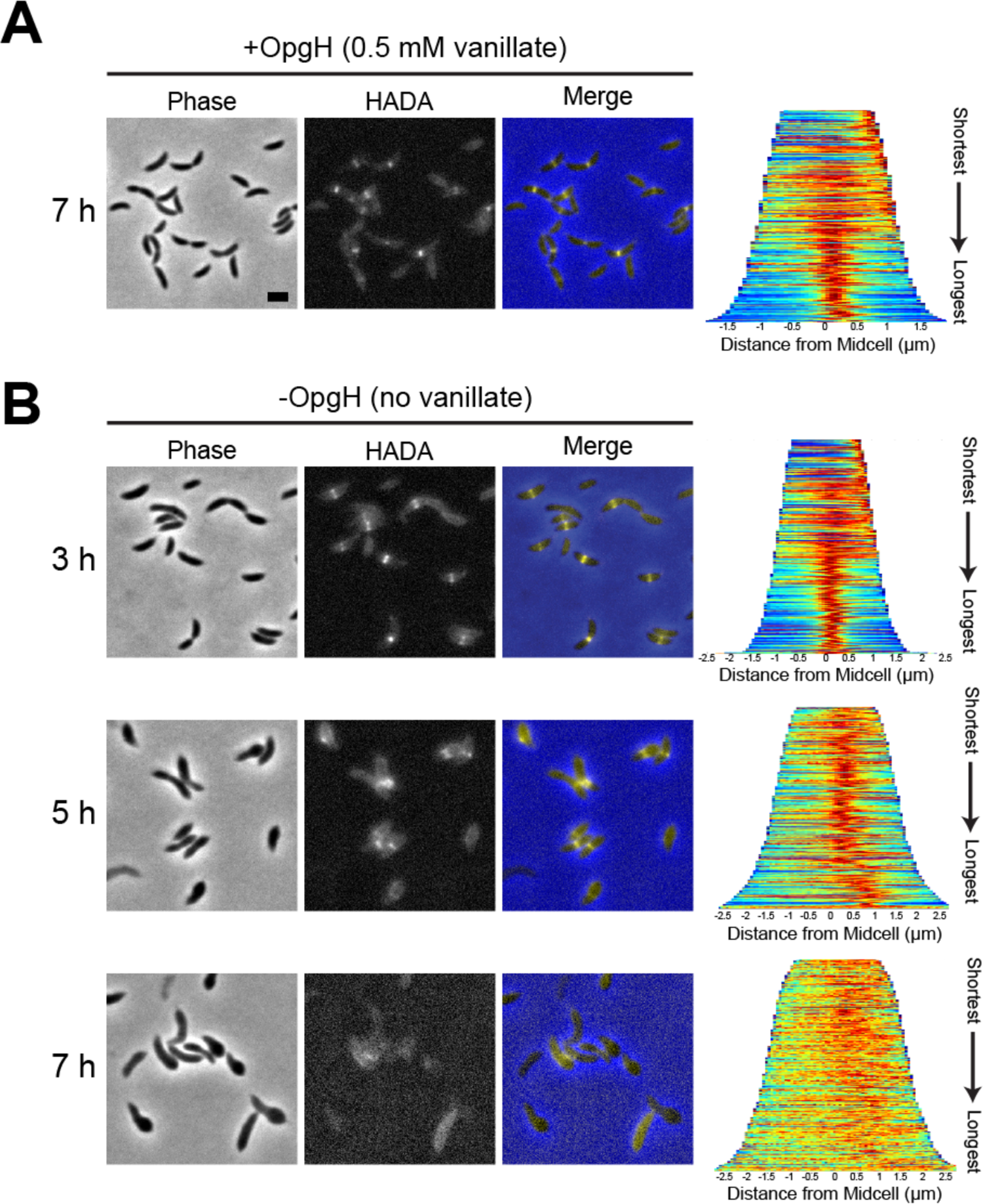
OpgH depletion results in misincorporation of cell wall material. Phase contrast, epifluorescence, and merged images showing HADA incorporation **(A)** with vanillate at 7 hours and **(B)** without vanillate over the course of 7 hours in the OpgH depletion strain (EG3421). Demographs (right) show the normalized HADA intensities across the population, arranged from shortest to longest cell for 300 cells. Scale bar, 2 µm.

### The divisome and elongasome are mislocalized in OpgH-depleted cells

The spatial misregulation of PG synthesis we observed during OpgH depletion aligns with the profound morphological defects that occur without OpgH. We hypothesized that misregulation of PG synthesis could be attributed to mislocalization of the PG synthetic machinery associated with the divisome or elongasome. We began with the elongasome and characterized the localization of mNeonGreen-tagged RodZ (mNG-RodZ) under the control of its native promoter. RodZ is an essential part of the elongasome complex and its localization corresponds to sites of PG synthesis (14). RodZ localization is dependent on the localization of MreB, the actin homolog that directs localization of elongasome proteins. In WT cells, RodZ and MreB are reported to exhibit patchy localization that focuses at midcell in stalked and predivisional cells (14, 15). We observed this expected localization pattern for mNG-RodZ in our OpgH depletion strain in the presence of vanillate (+OpgH) (Figure 5A). In contrast, when OpgH was depleted, RodZ became diffuse and created intense puncta along the cell by five hours of depletion, which was exacerbated by seven hours (Figure 5A). Additionally, we noticed an increase in mNG-RodZ signal over the course of depletion. Since *mNG-rodZ* expression was under the control of the native *rodZ* promoter, this could be attributed to an inability to turn over RodZ or a mechanism to increase production of RodZ as OpgH is depleted.

**Figure 5.**
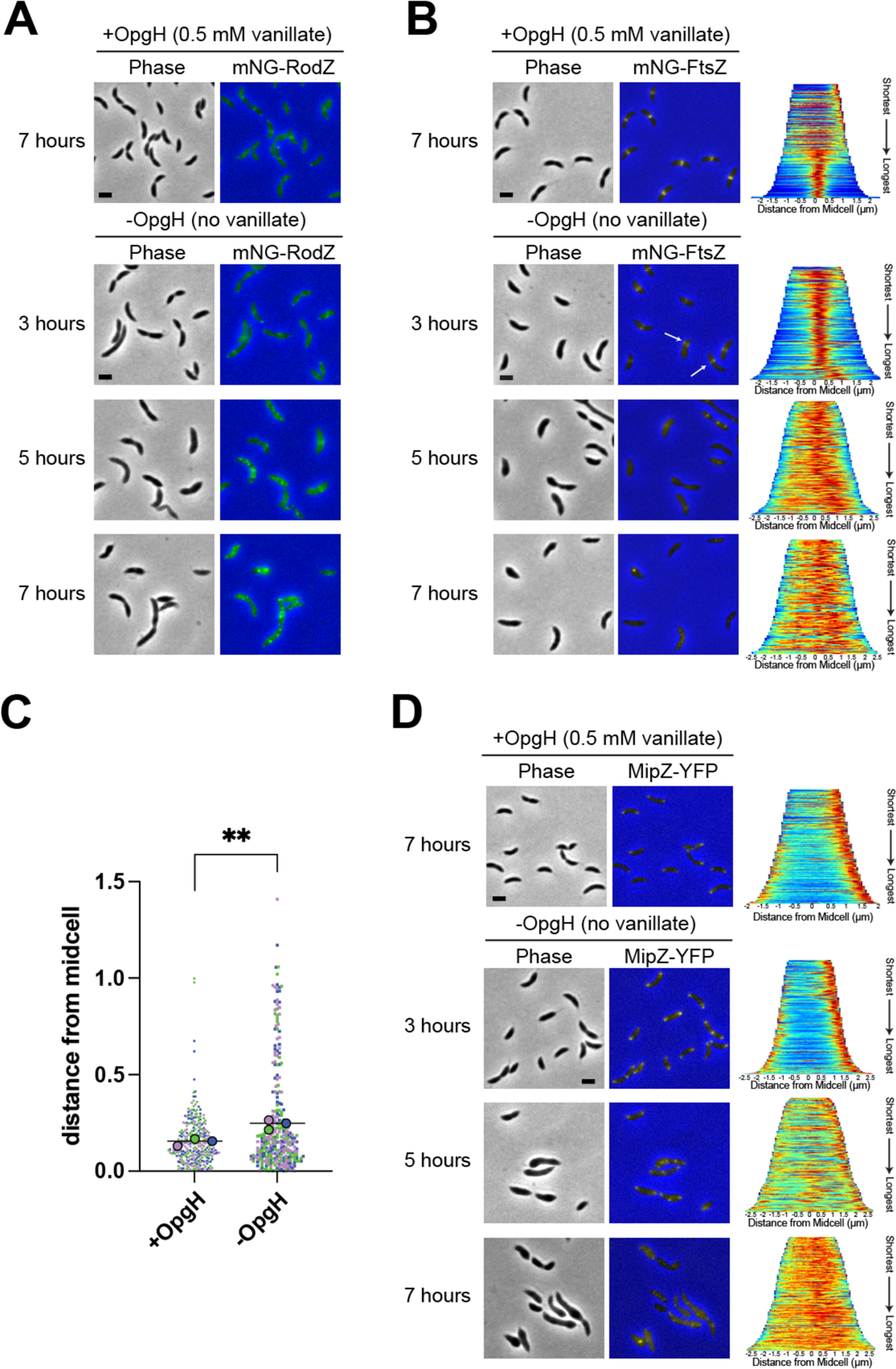
Divisome and elongasome proteins are mislocalized in OpgH-depleted cells. Phase contrast and epifluorescence merged images of **(A)** mNG-RodZ (EG3790), **(B)** mNG-FtsZ (EG3770), and **(D)** MipZ-YFP (EG3808) localization of OpgH depletion strains with (7 hours) and without 0.5 mM vanillate (at 3, 5, and 7 hours). White arrows indicate examples of asymmetric mNG-FtsZ localization. Demographs for FtsZ and MipZ localizations present the normalized intensities for cells across the population, arranged from shortest to longest (right). (**C**) Distance of the Z-ring from midcell (µm) in the presence of vanillate (+OpgH) or after OpgH depletion for 3 hours (no vanillate, -OpgH). Colors indicate measurements from three biological replicates. Small symbols reflect individual cell measurements, large symbols indicate mean for each replicate. Line indicates overall mean. **, p = 0.0074 between the means of the three replicate means using unpaired t-test. Scale bar, 2 µm.

Since RodZ’s midcell localization is also dependent on FtsZ, we next assessed FtsZ localization in the OpgH depletion strain. FtsZ, the master regulator of the divisome, is an essential protein. FtsZ forms a dynamic scaffold, called the Z-ring, that directs assembly of the divisome to the future division site (16). When OpgH was present, mNG-FtsZ consistently formed tight midcell bands in stalked and predivisional cells (Figure 5B, +OpgH). In contrast, when OpgH was depleted, mNG-FtsZ created slightly diffuse bands by three hours of depletion that appeared to be shifted away from midcell (Figure 5B, -OpgH, white arrows). This off-center FtsZ localization potentially aligns with the asymmetric HADA incorporation (Figure 4B) and bulging we observed. We determined the position of Z-rings in cells producing OpgH or cells depleted of OpgH for 3 hours and found that Z-rings were, indeed, positioned more asymmetrically (i.e. further from midcell) during OpgH depletion than in the presence of OpgH (Figure 5C). By five hours of depletion, FtsZ was almost entirely diffuse (Figure 5B). This aberrant Z-ring placement explains the inability of RodZ to properly localize, and suggests that, by five hours of OpgH depletion, both the divisome and elongasome are unable to properly localize and direct PG synthesis.

We next turned our attention to the Z-ring positioning protein, MipZ, which is a negative regulator of FtsZ assembly (17). MipZ follows the *Caulobacter* chromosome segregation machinery, forming a unipolar focus in swarmer cells, then becoming bipolar as the origin of replication is segregated. Bipolar MipZ inhibits FtsZ polymerization at the poles and directs Z-ring formation at midcell. MipZ-YFP localization in cells with OpgH was consistent with the previously reported unipolar to bipolar MipZ localization over the *Caulobacter* cell cycle (Figure 5D, +OpgH). After five hours of OpgH depletion, however, MipZ-YFP localization was largely perturbed, with more diffuse MipZ and with some cells exhibiting three or more MipZ foci (Figure 5D, -OpgH). After seven hours of OpgH depletion, MipZ-YFP was almost entirely diffuse, while only sometimes forming puncta (Figure 5D). We propose that mislocalization of MipZ in cells lacking OpgH is a key factor that leads to mislocalization of the divisome and elongasome, yielding the pleiotropic morphological defects we have observed. However, as FtsZ begins to mislocalize (at 3 hours post-depletion) prior to obvious disruption of MipZ localization (at 5 hours post-depletion) during OpgH depletion, multiple mechanisms may contribute to mislocalization of the morphogenetic machinery.

### The predicted structures of *Caulobacter* and *E. coli* OpgH suggest similarities and differences between OpgH in these species

OpgH has been best studied in *E. coli*, where it has been genetically and biochemically implicated in production of OPGs using UDP-glucose as a substrate (18, 19). *E. coli* OpgH (EcOpgH) has an auxiliary role, as well, acting as a direct inhibitor of FtsZ that coordinates cell size with nutrient availability (11). Specifically, deletion of *opgH* results in a mild reduction in cell length and overproduction of OpgH causes filamentation. The enzymatic activity of EcOpgH is not required to regulate FtsZ, and the FtsZ regulatory function has instead been attributed to the N-terminal region of EcOpgH. Since we observe morphological defects with depletion of *Caulobacter* OpgH (CcOpgH), we sought to compare the sequences and predicted structures of EcOpgH and CcOpgH to understand if CcOpgH may perform a similar moonlighting function to EcOpgH.

The Alphafold Protein Structure Database includes predicted structures for both EcOpgH (AF-P62517-F1) and CcOpgH (AF-B8GX72-F1), with very high model confidence scores for the majority of residues in each (20, 21). We aligned the primary sequences of EcOpgH and CcOpgH (Supplemental Figure 4A), as well as the predicted structures of each (Supplemental Figure 4B-D). The aligned structures have an overall rmsd of 0.678, indicating high structural similarity (Supplemental Figure 4D). The structures align particularly well in the predicted transmembrane regions and in the adjacent cytoplasmic domain predicted to be required for binding UDP-glucose, which is consistent with conservation of enzymatic function. However, as is clear from both the sequence and structural alignments, EcOpgH bears extensions in both the N-(gray) and C-terminal (salmon) regions compared to CcOpgH that fold into an extension of the cytoplasmic domain that is absent in CcOpgH (Supplemental Figure 4A,C). Moreover, there is little to no sequence conservation between the N- or C-terminal regions of CcOpgH and those of EcOpgH. Since FtsZ inhibition is attributable to the N-terminal region of EcOpgH, this, along with our observation that CcOpgH does not localize to midcell in *Caulobacter* cells (Supplemental Figure 3) and the distinct phenotypes observed with loss of OpgH between *E. coli* and *Caulobacter*, all suggests that CcOpgH likely does not regulate FtsZ in the same way as EcOpgH. We also note extensions in the predicted periplasmic loops of EcOpgH that are absent in CcOpgH. These could be sites where EcOpgH interfaces with periplasmic partners like OpgG, which is not encoded in *Caulobacter*.

### Glycosyltransferase activity of OpgH is required to maintain proper morphology

Given that OpgH is characterized as an OPG synthase in *E. coli* and the similarity between the predicted structures of *E. coli* and *Caulobacter* OpgH, we wondered if it has a similar enzymatic function in *Caulobacter*. We attempted to demonstrate a role for *Caulobacter* OpgH in producing OPGs using established methods for isolation and detection of *E. coli* OPGs (22). However, we were unable to detect OPGs using these methods, perhaps reflecting unique chemistry of *Caulobacter* OPGs compared to *E. coli* OPGs (7). Instead of assessing effects of OpgH on its putative product, therefore, we sought to test if depletion of OpgH altered cellular levels of its predicted substrate, UDP-glucose. To this end, we extracted and quantified polar metabolites from cells producing OpgH or cells depleted of OpgH for 5 hours in either defined (M2G) or complex (PYE) media. We found that levels of UDP-glucose increased 2- to 3-fold when OpgH was depleted in either media condition (Figure 6A, Supplemental Table 1), consistent with the hypothesis that *Caulobacter* OpgH converts UDP-glucose to an OPG product.

**Figure 6.**
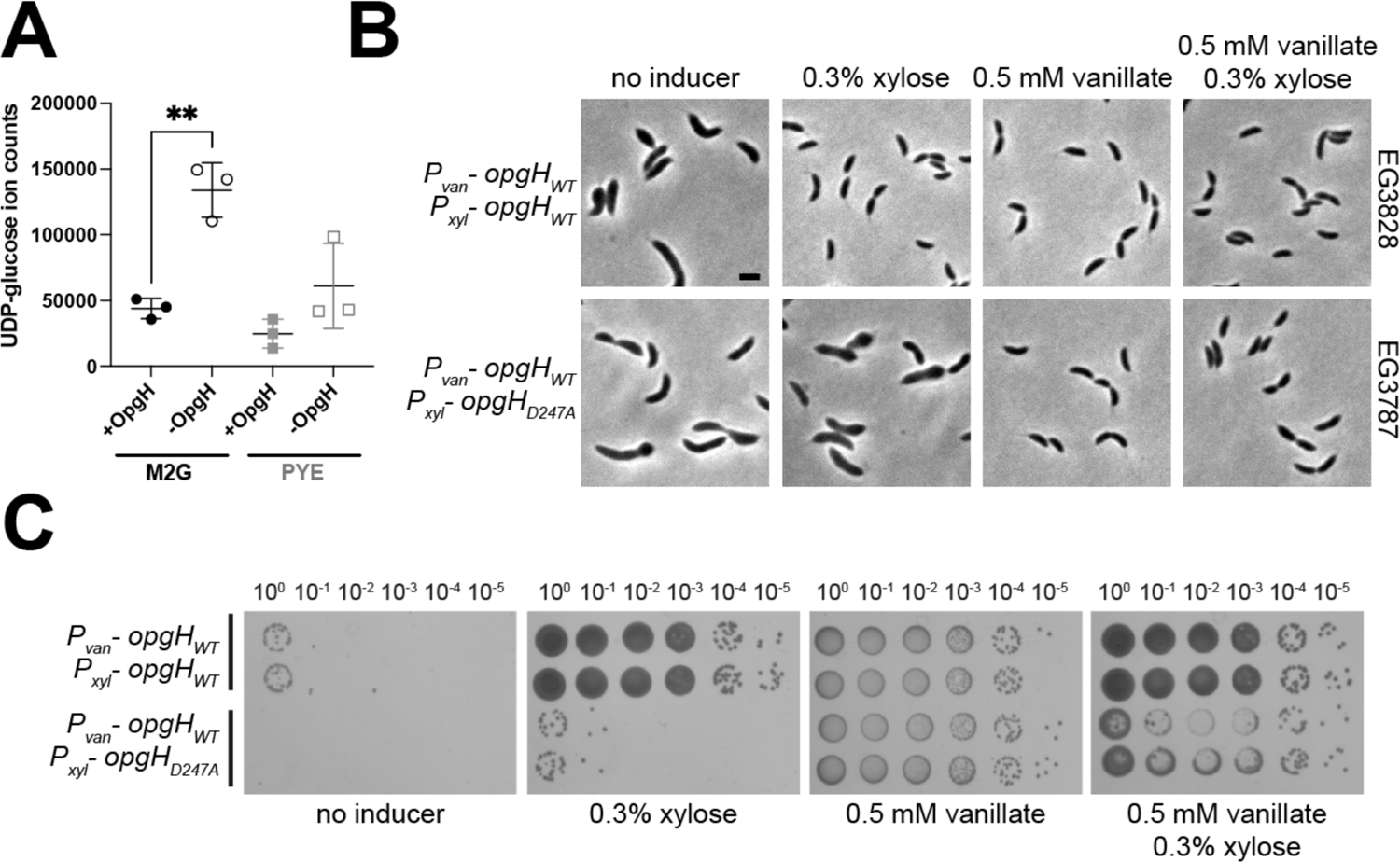
OpgH enzymatic activity is essential for morphogenesis. **A.** Levels of UDP-glucose in extracts from cells producing OpgH (+OpgH) or depleted of OpgH for 5 h (-OpgH) in the indicated media. Mean and standard deviation are plotted. **, p = 0.0022 using unpaired t-test. **B.** Phase contrast images of the OpgH depletion strain (Δ*opgH* + P_van_-*opgH*) with xylose inducible WT *opgH* (EG3828) or *opgH_D247A_* (EG3787). Cells have been depleted or induced with the indicated additive for five hours. Scale bar, 2 µm. **C.** Spot dilutions of the same strains grown on PYE with indicated inducer for two days. Two biological replicates of each are presented.

In *E. coli*, enzymatic activity of OpgH is dispensable for viability and for its effects on cell division. To probe the importance of the enzymatic activity of *Caulobacter* OpgH in promoting morphogenesis, we sought to characterize the role of predicted active site residues. Enzymes in the GT-A family of glycosyltransferases, like OpgH, all contain a conserved D-X-D motif (amino acids 245-247 (D-A-D) in *Caulobacter* OpgH) (Supplemental Figure 4A), which are responsible for binding the phosphate group on a nucleotide donor and coordinating a divalent cation required for activity (23). Mutation of either aspartate residue in the D-X-D motif eliminates the glycosyltransferase activity *in vitro* of other enzymes within this family, while not disrupting the fold of the enzyme (24).

We attempted to convert the latter aspartate in the *Caulobacter* OpgH D-X-D motif to an alanine (OpgH_D247A_) at the native *opgH* locus, but were unsuccessful. This initial observation suggested that the enzymatic activity of OpgH may be essential. We therefore investigated the phenotype of cells producing this variant protein by creating a strain harboring a native deletion of *opgH*, along with a vanillate-inducible copy of WT OpgH and a xylose-inducible copy of either WT *opgH* or the *opgH_D247A_* variant. Depletion of both copies of OpgH resulted in the expected elongation and bulging phenotype, and expression of WT *opgH* yielded normal growth and morphology. However, production of OpgH_D247A_ phenocopied depletion of OpgH (Figure 6B). We verified the lethality of the OpgH_D247A_ mutant with spot dilutions (Figure 6C). To ensure that the D247A mutation did not destabilize OpgH, we assessed the function and production of 3X-Flag-tagged WT or OpgH_D247A_ during depletion of OpgH. Consistent with the untagged variants, WT OpgH-3X-Flag supported growth while OpgH_D247A_-3X-Flag did not, despite each protein being stably produced as confirmed by immunoblotting (Supplemental Figure 5). The identification of a catalytically dead mutant of OpgH that phenocopies its depletion implicates OpgH’s glycosyltransferase activity in maintaining cellular morphology.

## Discussion

In this work, we have highlighted the essential and previously undiscovered role of OpgH in *Caulobacter* morphogenesis. We have demonstrated that *opgH* is essential (Figure 1) and characterized the unique morphological defects associated with loss of OpgH. Without OpgH, *Caulobacter* cells become elongated and develop asymmetric bulges (Figure 2). Bulges always occur in the stalk-proximal region of the cell, and result in lysis (Figure 3). Using PG labeling probes, we have shown that these phenotypes result from misregulation of PG insertion (Figure 4), driven by disruption of the divisome and elongasome (Figure 5). Interestingly, we can attribute the morphological defects of OpgH depletion to the glycosyltransferase activity of OpgH, as a catalytically dead OpgH mutant phenocopies the depletion (Figure 6).

Our data implicate the glycosyltransferase activity of OpgH in the morphological defects associated with its loss. However, we do not yet have direct evidence that *Caulobacter* OpgH can synthesize OPGs. Indeed, although OpgH-dependent synthesis of OPGs has been observed in, for example, *E. coli* cell extracts, OpgH activity has not been reconstituted with purified protein *in vitro* from any organism (18). We recently identified a genetic interaction between OpgH and the novel OPG metabolic enzymes, EstG and BglX (7). In that study, we determined that the substrate of EstG is a cyclic hexamer of glucose, which is the first OPG identified in *Caulobacter* (7). Although our data support the hypothesis that OpgH converts UDP-glucose to OPGs, further biochemical studies are necessary to determine if this cyclic OPG is, in fact, the product of OpgH.

Our results indicate the importance of OPG synthesis for localizing the divisome and elongasome. MipZ is delocalized in OpgH-depleted cells, suggesting a mechanism for mislocalization of FtsZ and other morphogenetic proteins (17, 25). Alternatively, loss of OPG synthesis might disrupt cellular energetics through glucose or carbon metabolism, which could consequently impact the morphogenetic complexes. This is plausible since both MipZ and FtsZ require energy-rich nucleotides (ATP and GTP respectively) to fuel their localization and dynamics (17, 26). As discussed above, *E. coli* OpgH has a direct connection with the divisome as an inhibitor of FtsZ. Though *Caulobacter* OpgH apparently lacks this interaction domain and does not localize with the morphogenetic complexes, we cannot rule out the existence of an interaction between OpgH and components of the divisome, elongasome, or other factors. However, given the importance of OpgH catalytic activity, our data suggests an effect of OPGs, themselves, on morphogenetic complexes. Mapping the biochemical and genetic interactions between OpgH and proteins involved in cell shape, as well as cataloging metabolic consequences of OPG production or depletion remain important avenues for future work.

Of the phenotypes associated with OpgH depletion, we were most surprised by the asymmetric nature of the bulging phenotype. Typically, cell bulging is a consequence of misregulation of either the elongasome/Rod complex or the divisome. Although *Caulobacter* cells are inherently polarized, the bulging that results from misregulation of the elongasome in *Caulobacter* (e.g. depleting MreB) (15) is uniformly distributed along the cell length, resulting in lemon-shaped cells. Bulging from a divisome mutant (i.e. a variant of FtsZ lacking its disordered linker (ΔCTL)) (27, 28), results in envelope bulges only where ΔCTL is localized, i.e. near midcell. The OpgH depletion phenotype is unique in its asymmetry, with bulging primarily on the stalk-proximal side of the cell. We observed that the Z-ring was more asymmetrically positioned after 3 hours of OpgH depletion than in the presence of OpgH. We therefore propose that depletion of OpgH causes a shift in the location of the divisome and elongasome away from the stalked pole at early stages of OpgH depletion. This would deplete PG synthesis from the stalk-proximal region of the cell and, if accompanied by ongoing cell wall hydrolysis in that region, lead to local cell bulging. Alternatively, or in addition, it is possible that the asymmetry in bulging is due to an unidentified factor that directs PG synthesis or hydrolysis asymmetrically. For instance, loss of OpgH might misregulate PG enzymes like L,D-transpeptidases, which primarily crosslink PG in the stalk (29, 30).

We have established that OpgH, and likely OPG production, plays a crucial role in *Caulobacter* morphogenesis. The essentiality of *Caulobacter* OpgH and morphological defects associated with its loss provide an opportunity to elucidate the mechanism of action of OpgH and the OPG biosynthesis pathway. The previously studied OpgH homologs have all been nonessential, which limits the questions that we can ask. For instance, it is more challenging to study functional mutants *in vivo* or conduct genetic screens in an organism where OpgH is nonessential. Thus, this provides an appealing possibility for future work on *Caulobacter* OpgH, including avenues such as a mutagenesis screen to isolate novel mutants or a larger scale functional domain analysis study. We have already identified functional OpgH mutants that suppress the lethality of cell envelope stresses in a hypersensitive mutant (7). These mutants, as well as extragenic mutations isolated from suppressor screens, will be valuable in elucidating the mechanistic role of OpgH and OPGs in cell envelope homeostasis. Our findings have established a fundamental homeostatic role for an essential OpgH homolog and have uncovered a novel connection between the OPG pathway and cellular morphology that is ripe for future investigation.

## Methods

### *Caulobacter crescentus* growth media and conditions

*C. crescentus* NA1000 cells were grown at 30°C in peptone-yeast extract (PYE) medium. Xylose and glucose were used at a concentration of 0.3% (w/v) and vanillate at 0.5 mM for induction/depletion experiments. Antibiotics used in liquid (solid) medium as are follows: gentamycin, 1 (5) µg/mL; kanamycin, 5 (25) µg/mL; spectinomycin, 25 (100) µg/mL. streptomycin was used at 5 µg/mL in solid medium. For growth curves, a Tecan Infinite M200 Pro plate reader measured absorbance every 30 minutes at OD_600_ of a 100 µL culture volume in a 96-well plate in biological triplicate with intermittent shaking. For spot dilutions, cells were grown to mid-log phase and diluted to an OD_600_ of 0.05. Cells were then serially diluted up to 10^-6^ and 5 µL of each dilution was spotted onto a PYE plate with indicated inducer and/or antibiotic. Plates were incubated at 30°C for 48 hours. Strains and plasmids used in this study are listed in Supplemental Table 2.

### Atypical strain construction

We could not make the following strains in low osmolarity PYE media, so they were constructed in M2G minimal media: EG3421 (*opgH::ΔopgH; vanA::P_van_-opgH*), EG3770 (*opgH::ΔopgH; vanA::P_van_-opgH*, *xylX::P_xyl_-mNG-FtsZ*), EG3790 (*opgH::ΔopgH; vanA::P_van_-opgH*, *xylX::P_rodZ_-mNG-rodZ)*, EG3808 (*opgH::ΔopgH; vanA::P_van_-opgH*, *xylX::P_xyl_-mipZ-YFP),* EG3828 (*opgH::ΔopgH; vanA::P_van_-opgH; xylX::P_xyl_-opgH*), EG3787 (*opgH::ΔopgH; vanA::P_van_-opgH; xylX::P_xyl_-opgH_D247A_*), EG3957 (*opgH::ΔopgH; vanA::P_van_-opgH; xylX::P_xyl_-3X-Flag-opgH*), and EG3959 (*opgH::ΔopgH; vanA::P_van_-opgH; xylX::P_xyl_-3X-Flag-opgH_D247A_*). For 500 mL of M2G plates, 465 mL of water and 7.5 g agar (1.5%) were autoclaved. Once cooled, 25 mL of 20X M2 salts, 500 µL of 500 mM MgSO_4_, 500 µL of 10 mM FeSO_4_ 10 mM EDTA (Sigma F-0518), and 0.3% glucose were added. Additional antibiotics or media supplements for selection were added at this time. After initial strain construction, all strains were able to be grown in either M2G or PYE liquid media with appropriate inducers.

### Microscopy

Cells in exponential phase were immobilized on 1% agarose pads and imaged using a Nikon Eclipse Ti inverted microscope equipped with a Nikon Plan Fluor 100X (NA1.30) oil Ph3 objective and Photometrics CoolSNAP HQ^2^ cooled CCD camera. Images were processed using Adobe Photoshop. Levels were adjusted to the same level across samples in a given experiment. Cell shape analysis was performed using CellTool and demographs were generated using Oufti. Z-ring position analysis was performed in MicrobeJ (31) using the “line” feature. Three independent replicates were imaged and analyzed for Z-ring positioning, and data were presented in SuperPlots (32). Inducible fluorescent fusion proteins were induced for 1 hour with 0.3% xylose before the indicated imaging timepoint.

### Metabolomics sample preparation and analysis

Metabolomic samples were prepared as described previously (33). Briefly, cells were grown to an OD600 of 0.3 in 4 mL and filtered through 0.22 μm nylon filters (Millipore GNWP04700). The cells were quenched by placing the filter upside down in a 60 mm dish containing 1.2 mL pre-chilled quenching solution (40:40:20 Acetonitrile:methanol:H2O + 0.5% formic acid) and incubated for 15 minutes at −20°C Cells were removed by pipetting the quenching solution over the filter, added to chilled bead beating tubes containing 50 mg of 0.1 mm glass beads, and neutralized with 100 μL 1.9M NH4HCO. Cells were lysed using a Qiagen Tissulyzer at 30Hz for 5 minutes. Samples were spun at 4°C for 5 minutes at 16,000 x g and transferred to pre-chilled tubes to remove debris.

Metabolomics was performed at the Metabolomics Core Facility at Rutgers-Robert Wood Johnson Medical School. Analysis used a Q Exactive PLUS Hybrid Quadrupole-Orbitrap mass spectrometer (Thermo Fisher Scientific) using hydrophilic interaction chromatography. LC separation included the Dionex UltiMate 3000 UHPLC system (Thermo Fisher Scientific) with XBridge BEH amide column (Waters, Milford, MA) and XP VanGuard Cartridge (Waters, Milford, MA). LC gradients were as follows: solvent A (95%:5% H2O:acetonitrile with 20 mM ammonium acetate, 20 mM ammonium hydroxide, pH 9.4); solvent B (20%:80% H2O:acetonitrile with 20 mM ammonium acetate, 20 mM ammonium hydroxide, pH 9.4); solvent B percentages over time: 0 min, 100%: 3 min, 100%; 3.2 min, 90%; 6.2 min, 90%; 6.5 min, 80%; 10.5 min, 80%; 10.7 min, 70%; 13.5 min, 70%; 13.7 min, 45%; 16 min, 45%; 16.5 min, 100%. Flow rate was 300 μL/min and injection volume 5 μL and temperature maintained at 25°C. MS scans used negative ion mode, resolution of 70,000 at m/z 200, and an automatic gain control target of 3 × 10^6 and scan range of 72 to 1000. MAVEN software package was used to analyze metabolite data (34).

### Immunoblotting

Western blots were performed using standard lab procedures. Log phase cultures were lysed in SDS-PAGE loading buffer and boiled for 10 minutes. Equivalent OD units of cell lysate were loaded. Standard procedures were followed for SDS-PAGE and protein transfer to nitrocellulose membrane. Antibodies were used at the following concentrations: Primary antibodies used were Flag-M2 - 1:1,000 (Sigma, St. Louis, Missouri) and CdnL −1:2,500 (33). Secondary antibodies used were 1:10,000 of HRP-labeled α-mouse (for Flag) (PerkinElmer) or α-rabbit (PerkinElmer) (for CdnL). Chemiluminescent substrate (PerkinElmer) was added to visualize proteins via an Amersham Imager 600 RGB gel and membrane imager (GE).

## Acknowledgements

We would like to thank the members of the Goley lab for helpful discussions and input. We also thank Joshua García Colón for his help on the project and Piyusha Mongia and Erika Smith for cloning assistance. We thank Jean Marie Lacroix for helpful discussions regarding OPGs. We thank Yujue Eric Wang and the Metabolomics Core Facility at Rutgers-Robert Wood Johnson Medical School for Metabolomics Core services. This work is funded by the NIH, National Institute of General Medical Science through R35GM136221 (E.D.G.) and T32GM007445 (training grant support of A.K.D.).

---

**Supplemental Figure 1.**
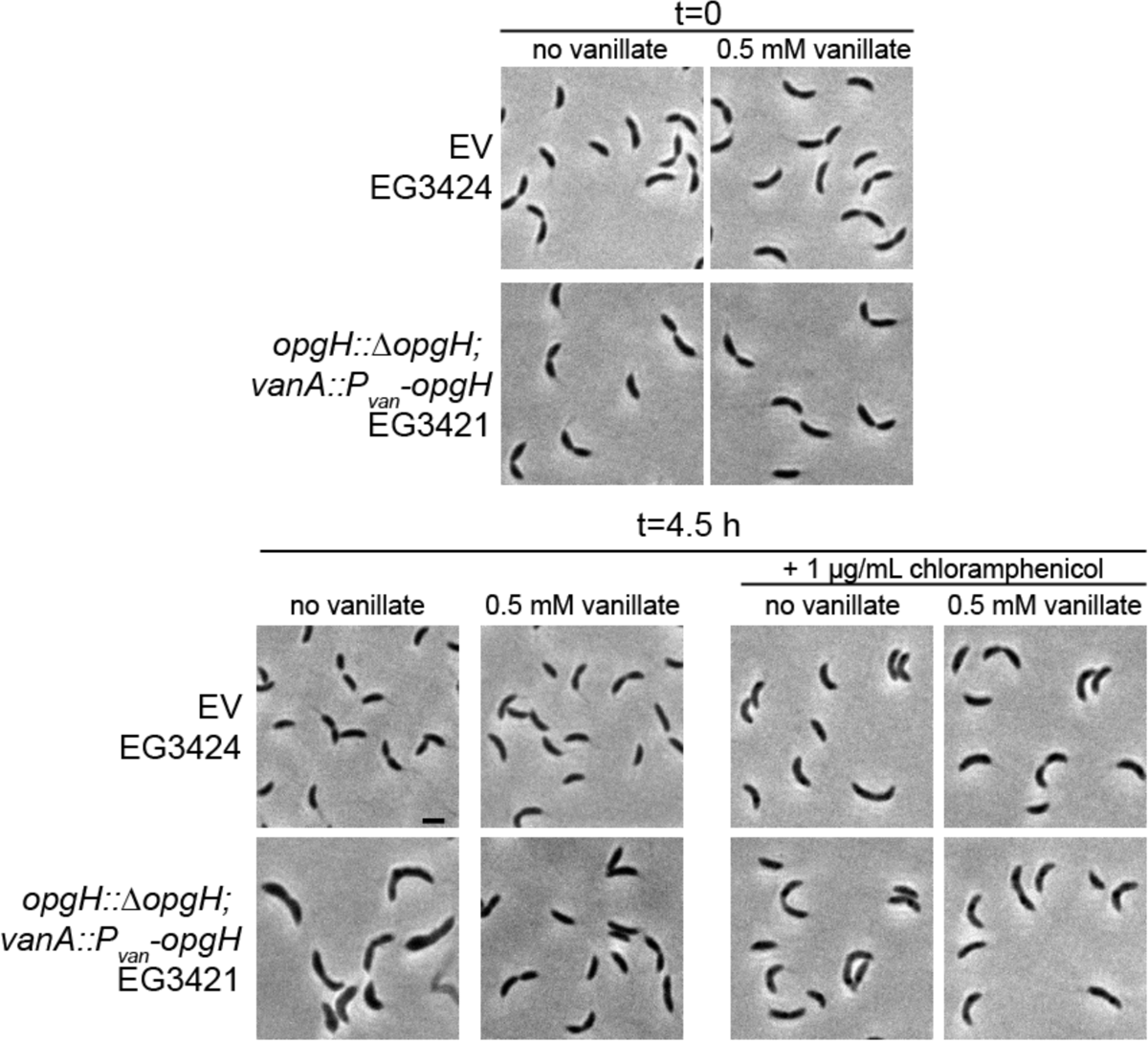
The OpgH depletion phenotype is reliant on cell growth. Phase contrast images of and empty vector (EV, EG3424) and OpgH depletion (EG3421) strains at the start (t=0) and end (t=4.5 hours) of treatment with sub-lethal chloramphenicol (1 µg/mL) in the presence and absence of vanillate. Scale bar, 2 µm.

**Supplemental Figure 2.**
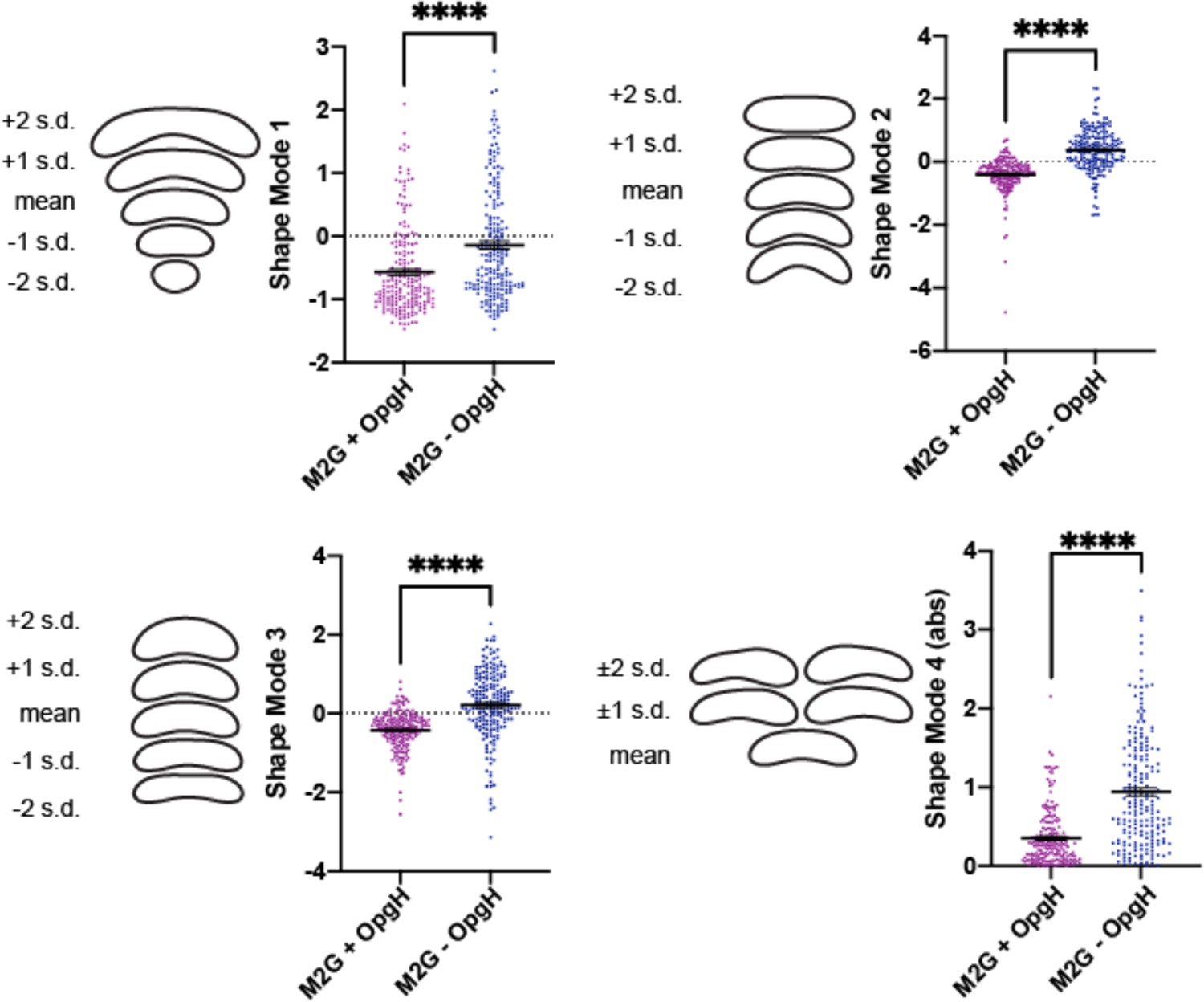
OpgH depletion causes morphological defects in minimal media Principal component analysis (PCA) of the OpgH depletion strain (EG3421) after 5 hours grown in M2G with 0.5 mM vanillate (purple, +OpgH) or without vanillate (blue, -OpgH). Scatter plots of 200 cells are presented. Shape modes 1, 2, 3, and 4 correspond to length, curvature, width, and asymmetric bulging. Contours indicate the mean shape and 1 or 2 standard deviations from the mean. Shape mode 4 shows the absolute value. Statistical analysis uses a Mann-Whitney unpaired t-test. **** = P < 0.0001.

**Supplemental Figure 3.**
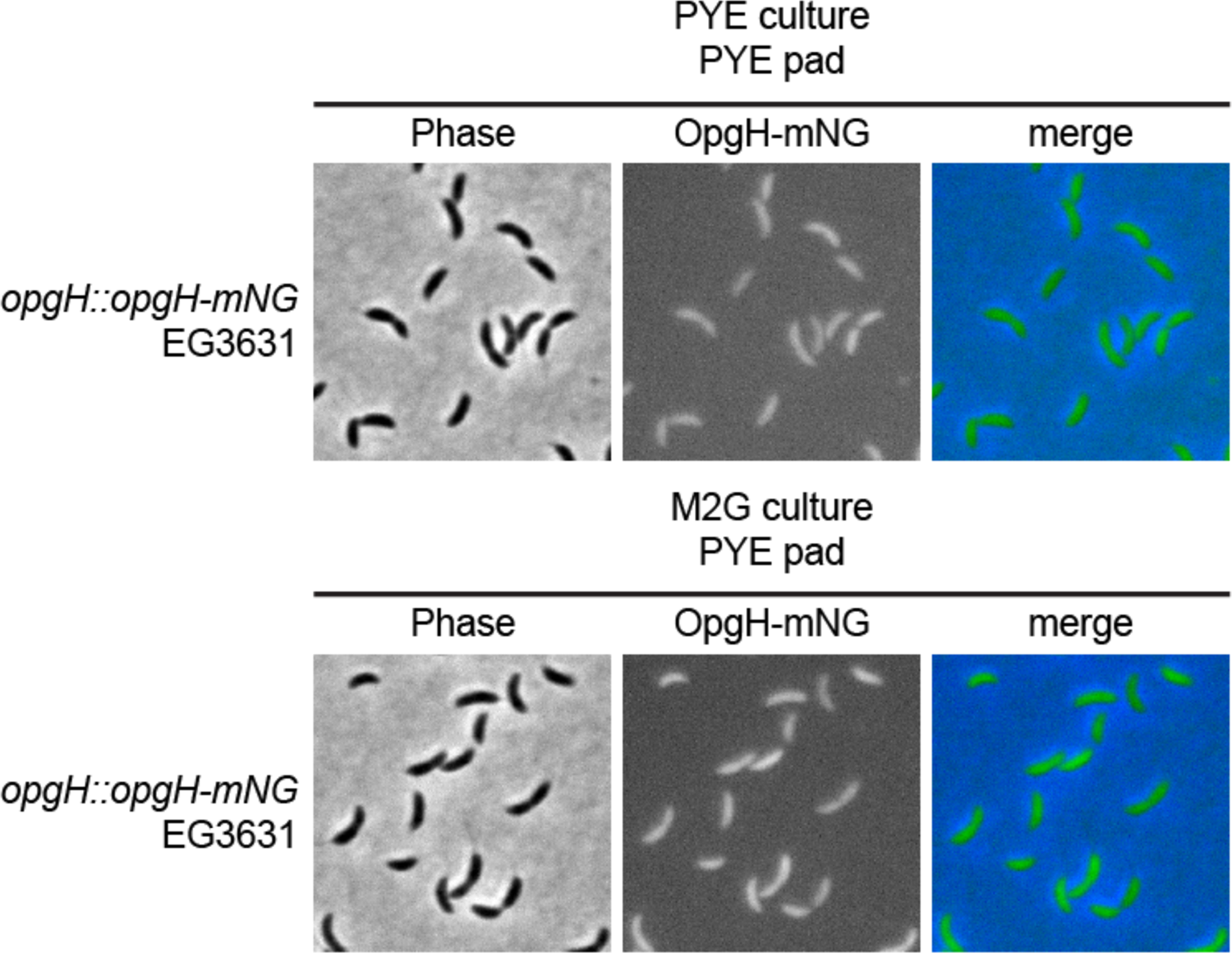
OpgH is diffuse within the cell. Phase contrast, epifluorescence, and merged images showing localization of OpgH with a C-terminal mNeonGreen (mNG) tag (EG3631) grown in PYE or M2G.

**Supplemental Figure 4.**
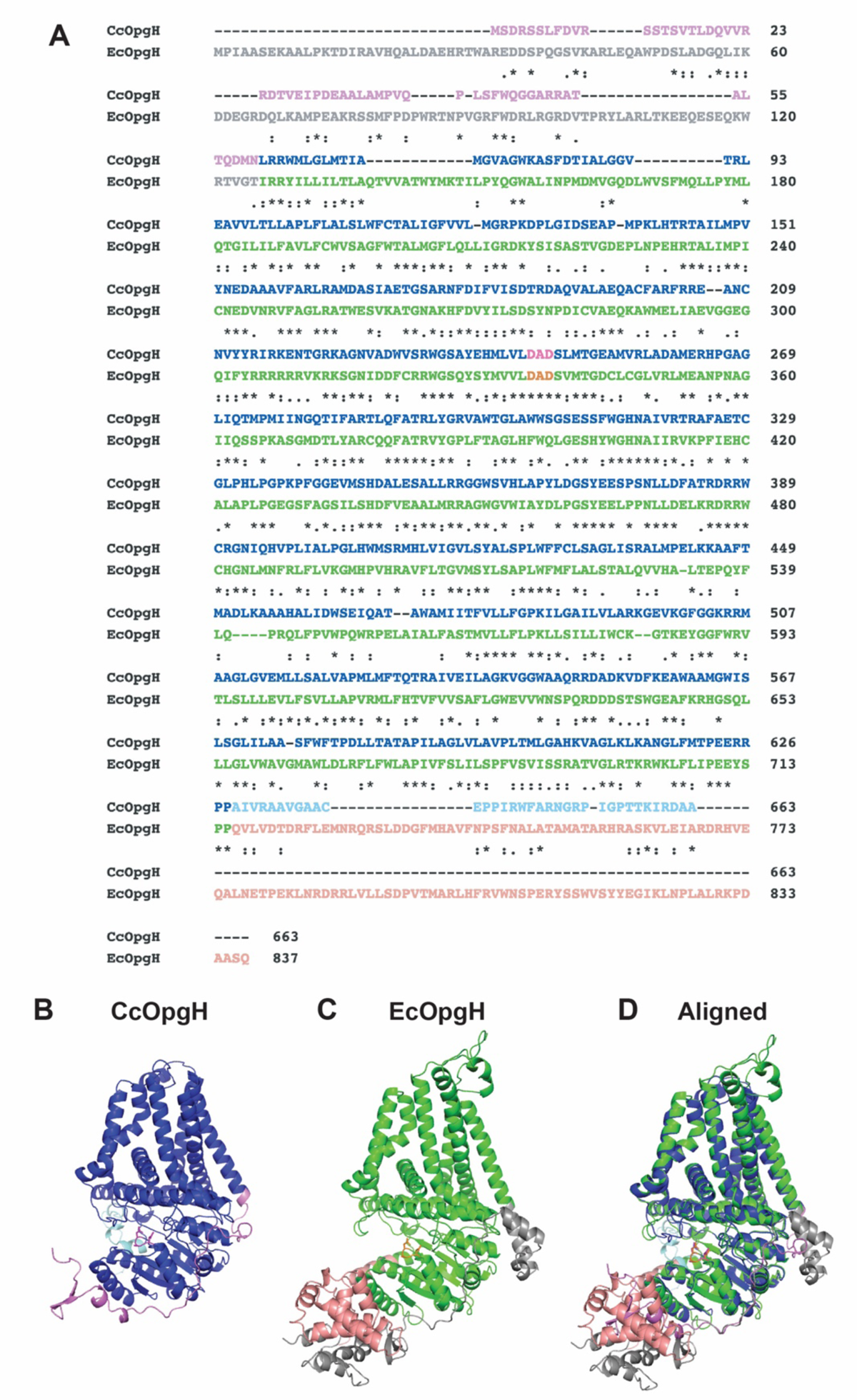
Sequence and structure alignments of *E. coli* and *Caulobacter* OpgH reveal similarities and differences. **A**. The primary sequences of *Caulobacter* (CcOpgH) and *E. coli* (EcOpgH) were aligned Clustal Omega. The poorly conserved N- and C-terminal regions of each are colored independently from the well-conserved central region. The predicted catalytic D-A-D residues of each are highlighted in magenta (CcOpgH) and orange (EcOpgH). **B** and **C**. Alphafold predicted structures of CcOpgH (B) and EcOpgH (C) with residues colored as in (A) and oriented with the predicted periplasmic face at the top and cytoplasmic regions at the bottom. **D.** The structures in B and C were aligned in Pymol yielding an rmsd of 0.678. Major differences observed are in the N- and C-termini, as well as in predicted periplasmic loops of EcOpgH.

**Supplemental Figure 5.**
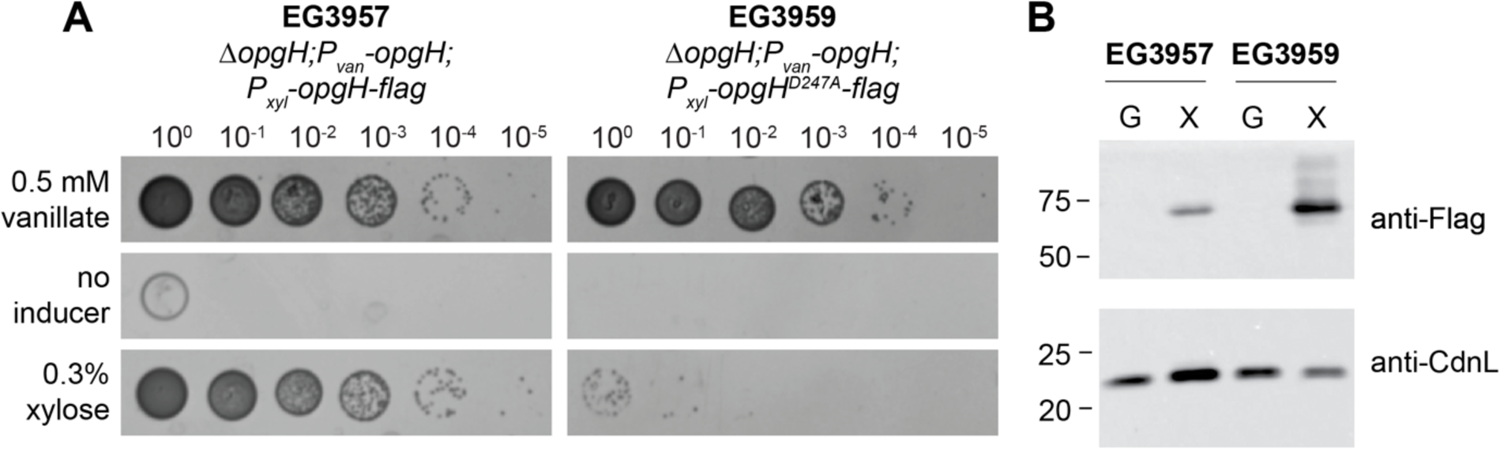
3X-Flag-tagged OpgH_D247A_ cannot complement loss of OpgH but is stably produced. **A.** Spot dilutions of the OpgH depletion strain (Δ*opgH* + P_van_-*opgH*) with xylose inducible WT *opgH-3X-Flag* (EG3957) or *opgH_D247A_-3X-Flag* (EG3959) grown on M2G with indicated inducer for two days. **B.** Immunoblot of lysates from the indicated strains grown in PYE with glucose (G) or xylose (X) for 6.5 hours. CdnL was used as a loading control.

**Supplemental Table 1.** Polar metabolites in extracts from cells producing OpgH or depleted of OpgH for 5 h in M2G or PYE. Metabolites reduced at least two-fold during OpgH depletion in both media conditions are highlighted in red, those increased at least two-fold during OpgH depletion in both media conditions are highlighted in green.

**Supplemental Table 2.** Strains and plasmids used in this study.

## Notes

### Competing Interest Statement

The authors have declared no competing interest.

